# Climatic limits to Atlantic salmon population fitness at continental scales

**DOI:** 10.1101/2023.12.18.571080

**Authors:** Olivia Morris, Hlynur Bárðarson, Alexia González-Ferreras, Rasmus Lauridsen, Samraat Pawar, James Rosindell, Guy Woodward

## Abstract

Anadromous fish populations are declining globally, partly due to acute pressure from rapid environmental change in their freshwater and marine habitats. A more mechanistic understanding of how climatic and land use changes impact their population-level fitness is needed to mitigate these declines. Here we develop a model that successfully captures the thermal niche of the declining Atlantic salmon. This allows us to predict the combined effects of changes in two dominant threats to this species: climate and resource availability. Specifically, the model predicts that a 50% reduction from metabolically optimal resource supply could constrict their thermal niche by ∼7°C. We also show that daily and seasonal temperature fluctuations have a relatively minor impact. A conservative increase of 1.5°C in global temperatures will cause fitness declines for populations in higher climatic regimes, across resource levels. Our results provide new and general insights into factors limiting the distribution of extant Atlantic salmon populations. They also highlight the relative importance of future threats from climatic warming, fluctuations, and changes in resource availability due to land use change.

---

Fish populations worldwide are facing rapid population declines, with many estimated to be at risk of local or global extinction^1–4^. In particular, populations of migratory fishes that partly or exclusively use freshwaters have declined by an average of 76% globally^3^. These include anadromous species, which live in the ocean as adults and migrate to freshwaters to reproduce. Atlantic salmon (*Salmo salar*) is an anadromous species that is particularly important for its ecological, cultural, and economic value^5^. Populations have declined by around 70% in the last 25 years, with extirpations from many rivers across its range^6–12^. Whilst there have been locally-effective efforts to mitigate these declines^13^, most populations continue to decline across the species’ distribution ^14,15^.

For Atlantic salmon and anadromous fish in general, environmental change in the form of combined climate and land use change is a pervasive threat to both the freshwater and marine phases of their complex life cycle^2,16,17^. During the marine adult stages of such species, there is growing evidence that climatic warming is already altering survival rates and dynamics of the wider food web^14,18–20^. Environmental changes during the juvenile freshwater stages of anadromous fish however, are of equal or greater concern due to the dominant contribution of these stages to population-level fitness in vertebrates ^21,22^. This is because temperature changes alter individual metabolism of these strongly ectothermic life stages, as well as their resource availability by changing primary productivity and prey abundance throughout the food web ^23–25^. In particular, warming increases metabolic demand, affecting juvenile development and mortality rates, ultimately changing population fitness ^22,26^. Furthermore, thermal fluctuations, which are expected to increase with climate change, are expected to depress fitness and narrow thermal niches^27–30^. These effects of temperature-driven metabolic perturbations may be further compounded by temperature-independent changes in resource inputs into freshwaters due to land use changes^31^. Thus, to predict how anadromous species such as Atlantic salmon will respond to a changing environment, we need a mechanistic understanding, rooted in physiology, of the climatic constraints on their freshwater life stages in particular, including its fundamental thermal niche^32–34^. That is, the range of temperatures over which a population can be viable (henceforth, just thermal niche). Despite this, to the best of our knowledge, there is no metabolically explicit framework to predict the thermal niche for migratory fish under novel environmental conditions arising from future climatic change and shifts in resource availability.

We developed a new mechanistic model for the thermal niche of Atlantic salmon: the temperature dependence of maximal population growth rate (*r*_max_) as a measure of density-independent fitness ^35–39^. To this end, we incorporated metabolic traits into an Integral Projection Model (IPM)^40^. The resulting Metabolic IPM (MIPM) combines the metabolic constraints of temperature and body size on the three key life-history traits—individual growth, mortality, and fecundity rates—across juvenile and adult stages to determine the population’s *r*_max_ (Fig. 1). Using the MIPM, we examined how the thermal niche changes through the combined effect of chronic warming, thermal fluctuations, and temperature-independent changes in resource availability on *r*_max_ of Atlantic salmon populations. We set out to examine the thermal niche and effects of warming and resource supply in Icelandic rivers and later expanded the scope to include a much broader latitudinal range. Our rationale is to use the mechanistic nature of our model to transfer understanding about the species in general between different geographic regions.

**Figure 1.**
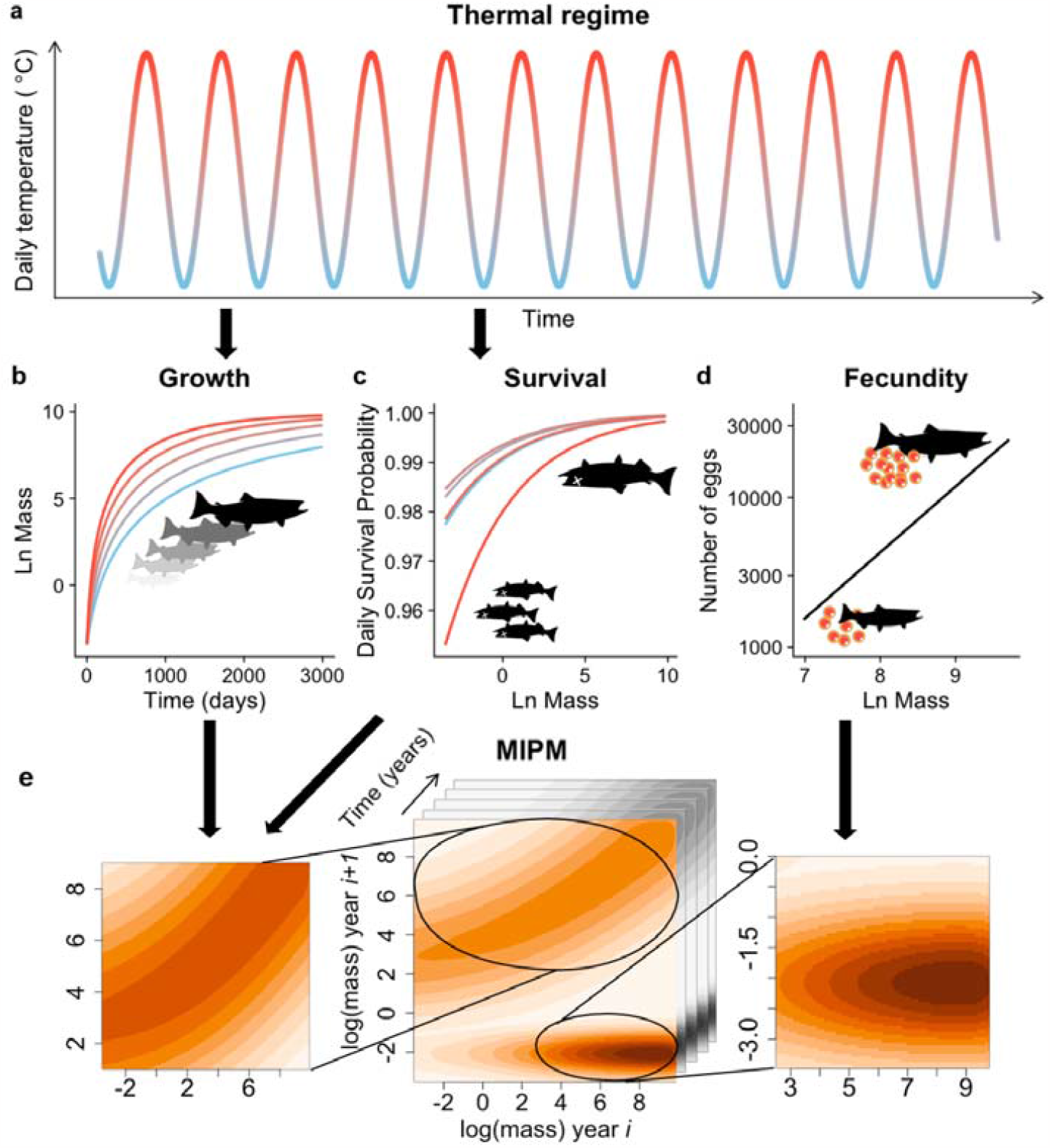
Illustration of the Metabolic Integral Projection Model (MIPM). The thermal regime (based on real time series data) (a) drives metabolic life-history traits (vital rates) as functions of body mass (grams) and temperature, for growth (b) and survival (c). These are used to parameterise a IPM kernel for the probability of surviving and growing across days, through successive life stages. At the onset of reproduction, fecundity (which scales with adult body mass; (d)), is added to the MIPM kernel as a single spawning event within a year (e). Daily thermal fluctuations are explicitly accounted for within a single ‘year matrix’. Note that small size classes have a disproportionate impact on the emergent maximal population growth rate (*r*_max_).

The MIPM predicts a left-skewed, unimodal form of the thermal performance curve (TPC) of *r*_max_ for Atlantic salmon (Fig. 2a), as expected for ectotherms in general^41–43^. To the best of our knowledge, this is the first ever characterisation of the TPC of fitness and the fundamental thermal niche for this species. The shape of this TPC remains qualitatively the same across all temperature and resource scenarios and arises from trade-offs between the underlying temperature-dependent traits: because individuals grow faster under warmer conditions, cumulative mortality across life stages decreases because mortality decreases with size. However, beyond the thermal optimum, mortality increases exponentially, outweighing the benefits of the faster development rate.

**Figure 2.**
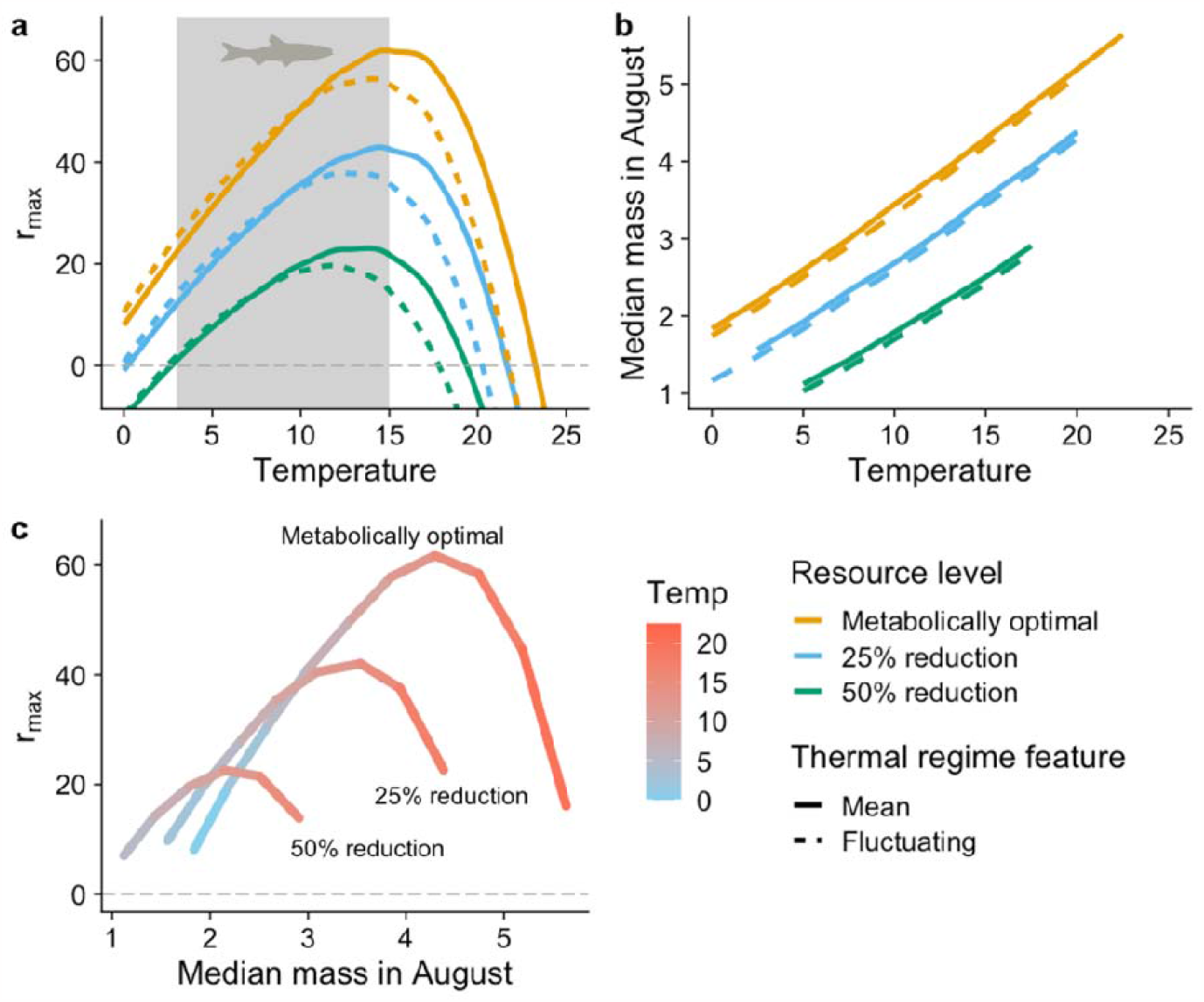
The effect of differing thermal regimes and resource level on the temperature-dependence of population fitness of Atlantic salmon and on the mean August size distribution. Increasing the mean temperature (horizontal axis) under mean (solid line) and fluctuating (dotted line) thermal regimes (°C) results in a unimodal relationship of *r*_max_ irrespective of resource level (a). The grey shaded area corresponds to the extant realised thermal niche of Atlantic salmon (see Methods). Note that in natural geographic gradients the amplitude of fluctuations around each population’s experienced mean temperature increases with latitude (change in thermal regime). (b) The effect of increasing the mean temperature under mean and fluctuating thermal regimes on the median mass calculated from the averaged August size distribution (log grams). Where *r*_max_ is greater than or equal to zero, increasing mean (solid lines) and fluctuating (dotted lines) thermal regimes increase the median mass during August from the mean size distribution for all resource levels. If *r*_max_< 0, the mean stable size distribution is not well defined. (c) *r*_max_ increases with the median of the population’s body mass distribution averaged for August to around 15°C before rapidly declining. This is mediated by resource level and exemplified with an increasing mean thermal regime.

The MIPM also predicts that at lower resource levels, the temperature at which *r*_max_ is maximised (thermal optimum) decreases, along with the thermal niche width (where, *r*_max_) (Fig. 2a), consistent with empirical studies on other ecotherms^26,44,45^. That is, warming creates a physiological deficit if the ecosystem does not have sufficient resources to support the increased pace of life^46^. Niche theory also supports this, as a reduction in resource supply is expected to constrict the thermal niche^33,47,48^. Upon overlaying the annual mean temperature range of freshwaters that extant Atlantic salmon inhabit (the realised thermal niche width), we find that current populations occupy a subset of their fundamental thermal niche at any given resource level (Fig. 2a). This is consistent with the fact that organisms typically buffer the risk of “falling off” their thermal fitness peak due to fluctuations^27,44,49^. Furthermore, it is noteworthy that the most restricted resource scenario (50% reduction) is the one that best captures the observed latitudinal distribution of extant Atlantic salmon (Fig. 2), consistent with the expectation that resource levels in terms of prey availability in the wild are typically low^50–53^.

The MIPM also shows that mediated by the resource regime, the median of the population’s body mass distribution across life stages (the size-stage structure; Supplementary Fig. 2) increases linearly with temperature (Fig. 2b). As a result, within the observed temperature range of Atlantic salmon populations, *r*_max_ increases with the median mass, also mediated by the resource regime (Fig. 2c). The MIPM thus provides a mechanistic explanation for the link between the size-stage distribution and population fitness, where larger body size is typically associated with greater fitness^54^. Beyond ∼15°C population fitness declines as the warmer temperatures which result in these larger masses have higher mortality. This has practical implications: because the local size distribution is relatively feasible to measure in the field^11^, this framework allows the fitness of field populations to be inferred from snapshots of the population’s size-stage distribution. This provides practitioners with a relatively simple quantitative tool to monitor and compare the levels of metabolic and fitness stress on populations across their geographical and latitudinal ranges.

We also investigated the importance of natural (seasonal) climatic fluctuations in temperature on the TPC of *r*_max_ in Atlantic salmon. In general, at any given resource level, although fluctuations lower the thermal optimum along with the critical thermal maximum (the maximum temperature where *r*_max_), overall, they have a relatively small impact compared to changes in mean temperatures across latitude (Fig. 2a).

Finally, we explored effects of the constraints imposed by the Atlantic salmon thermal niche on their ability to respond to ongoing and future climatic warming. The most conservative IPCC scenario of mean global warming of 1.5°C (SSP1-1.9^55^), is predicted to reduce fitness, especially if resource levels also decrease (Fig. 3a). A greater increase of 3.6°C (IPCC SSP3-7.0^55^), will cause further, more drastic declines in population fitness compounded by resource supply and thermal fluctuations (Fig. 3b). Thus, one way that Atlantic salmon populations can cope with environmental change is by increasing resource intake. Indeed, some recent studies suggest that salmonids can cope with the increasing metabolic demands of a warming climate by switching to alternative prey or increasing baseline consumption rate^56,57^. There is also evidence that adult Atlantic salmon are beginning to migrate further North when at sea in response to warming^58,59^. Whilst these strategies may be possible in some Atlantic salmon populations, our results highlight the fact that if prey availability is to remain constant, decrease or disconnect with warming as has been predicted in temperate freshwaters ^14,25,60,61^, warming will raise possibly insurmountable challenges in the path of their ability to adapt to climate change. Our results thus emphasise the importance of the freshwater stages, which cannot easily shift ranges in response to warming. Furthermore, as climatic change is also predicted to entail increased fluctuations in temperature and resources, this would have stronger negative impacts in the cooler (northern) regions of the species’ range where baseline fitness is already low (Fig. 2a).

**Figure 3.**
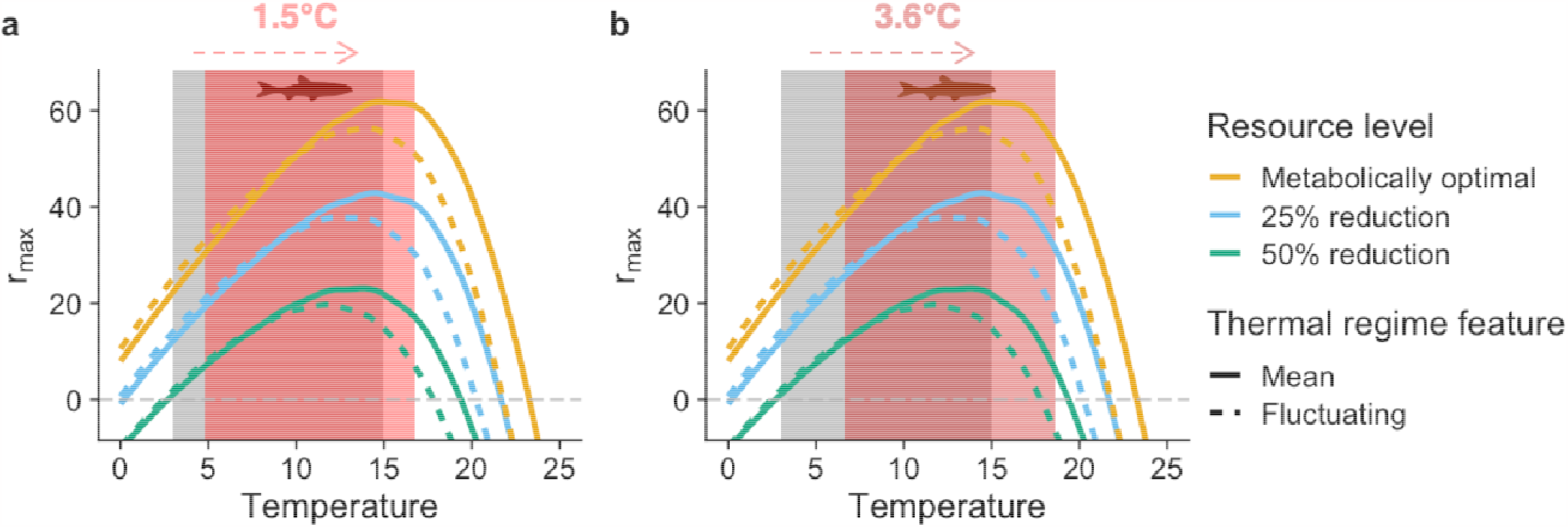
Mean annual temperature changes under two IPCC scenarios on the realised thermal niche of Atlantic salmon. Assuming Atlantic salmon remain within their current geographic distribution, the effect of a global mean increase in environmental temperature of (a) 1.5°C by 2050 (SSP1-1.9^55^) and (b) 3.6°C by 2100 (SSP3-7.0^55^), on the realised thermal niche and the maximal rate of population growth, *r*_max_. The current realised (empirically observed) thermal niche is shown in grey with the IPCC scenario overlaid in red. Note that under both global warming scenarios, overall population fitness (area under the curve) reduces.

These results suggest that mitigation against climate change could therefore be considered for Atlantic salmon populations by focusing on the upper range of the observed (realised) thermal niche within their freshwater stages. For example, riparian tree planting may increase carbon and nutrient supply available to these vulnerable life stages through the food web^62–64^. We recognise however, that the MIPM does not include the effect of physiologically explicit parameters such as oxygen supply. Warmer temperatures also increase oxygen demand, therefore warming will limit the aerobic scope and place a greater metabolic load^65,66^. This may cause the MIPM to underestimate the impact of warming as oxygen may therefore further limit performance and reduce population fitness at higher temperatures^44^. Our model has also assumed within species consistency in metabolic traits, it would be interesting to test this assumption with future work and to discover if, for example, local adaptation has occurred at thermal extremes.

Our study provides novel insights into the metabolic stress and fitness levels of Atlantic salmon populations across their extant geographical range as well as their potential to cope with future environmental changes. We hope that our MIPM will act as a key stepping stone towards developing the next generation of mechanistic models needed to conserve and manage global populations of Atlantic salmon in Iceland and elsewhere across their global range. Beyond this, we expect that the new MIPM framework will be applicable for predicting the impacts of climate change on other focal species where data on their temperature-dependence of fundamental life-history traits (growth, survival, and reproduction rates) are available.

## Methods

### The Metabolic IPM

We build on the existing Integral Projection Model (IPM) concept, which was designed to model population dynamics driven by continuously varying traits such as body size ^40^. In an IPM, the mass distribution of individuals at time *t* changes over a fixed discrete time step governed by the equation

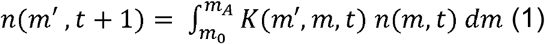

where *n* (*m,t*) is the number of females of body mass *m* (in grams) at time *t*, and *K*(*m*′, *m, t*) is a kernel capturing the growth and survival of Atlantic salmon of mass *m* into Atlantic salmon of mass *m*′ one discrete time step later. The equation is integrated between the minimum individual embryo mass, *m*_0_, and the maximum mass, or asymptotic mass, *m*_*A*_ (defined in Table 1). The kernel (*K*) is composed of two component kernels *P*, the growth and survival kernel, and *F*, which is related to fecundity. We can therefore write that *K*(*m*′, *m, t*) = *P*(*m*′, *m*,) + *F*(*m*′, *m, t*). The functions *P* and *F* can be further decomposed as

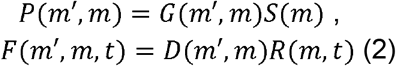

where *G*(*m*′, *m*) is the probability that an individual will grow from mass *m* to mass *m*′ over the course of a single time step and, subject to this, *S*(*m*) is the probability that an individual of mass *m* survives to the next time step. The *F*(*m*′, *m, t*)function within the kernel, includes *D*(*m*′, *m*,, which is the mass distribution of eggs and incorporates the probability that an egg produced by an individual of mass *m* are of mass *m*′, as well as the reproduction function, *R*(*m,t*), which is the number of eggs produced between time *t* and *t* + 1 by an individual of mass *m*. In cases of seasonal spawning events *R*(*m,t*)= 0 for most values of *t* and *R*(*m,t*) >0 specifically at the moments of spawning events.

**Table 1.** Metabolic parameters.

| Parameter | Description | Value | Unit | Reference |
| --- | --- | --- | --- | --- |
| $m$ | Body mass | - | g | - |
| $t$ | Time | - | day | - |
| $T_c$ | Temperature | - | °C | - |
| $m_0$ | Minimum mass of embryo | 0.03 | g | 101 |
| $m_A$ | Asymptotic mass | 20000 | g | 11,102 |
| $a(T_0)$ | Energy allocation rate | Metabolically optimal: 0.0146<br>25% reduction: 0.011<br>50% reduction: 0.007 | $\text{g}^{1/4} \text{d}^{-1}$ | 103–105 |
| $E$ | Activation energy | 0.45 | eV | 71 |
| $k$ | Boltzmann's constant | $8.62 \times 10^{-5}$ | $\text{eV} \cdot \text{K}^{-1}$ | |
| $T_0$ | Reference temperature | 273 | K | 106 |
| $z_0$ | Mortality scaling constant | 0.009 | | 82 |
| $E_D$ | Deactivation energy | 2.106 | | 82 |
| $T_{pk}$ | Peak performance temperature | 9.0579 | °C | 82 |
| $r_0$ | Fecundity normalisation constant | 1.303 | | 87–89 |
| $\beta$ | Fecundity scaling exponent | 1.013 | | 87–89 |
| $m_\alpha$ | Size at maturity | 1600 | g | 107 |

To parameterise the IPM, *G*(*m*′, *m*) and *S*(*m*) are derived from the Metabolic Theory of Ecology (MTE) and are independent of density. By constraining the IPM in this way, we circumvent the complexity of a more explicitly physiological model, for example Dynamic Energy Budget Theory parameterised models ^67,68^, but we still allow characteristic timescales and rates to be parameterised in a realistic way. The metabolic rate (rate of energy use) of an individual (*B*) can be expressed in terms of body mass and temperature as ^36^:

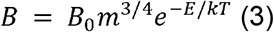

where *m* is the mean body mass (in our case, mean across individuals in a particular size/age class) and *T* is temperature in Kelvin, *k* is Boltzmann’s constant and *E* is the apparent activation energy, in eV, arising from all the metabolic reactions underlying the rate. *B*_0_ is a size- and temperature-independent normalisation constant that can vary with the environmental context or taxonomic group. An IPM’s vital rates of growth and mortality vary according to MTE laws, scaling as a function of body mass or temperature ^69–71^. Fecundity (*R*(*m,t*)) and egg mass distributions (*D*(*m*′, *m*)) are based on data from the literature relating these functions to Atlantic salmon body mass.

### Growth

To incorporate the effect of temperature on growth, we use MTE’s ontogenetic growth model ^72^:

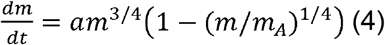

This model assumes that the rate of increase in mass is equal to the rate of acquisition of nutrients and energy by a growing individual, minus the cost of maintenance. More precisely, the rate of change of mass (*m*) with respect to time (*t*) is expressed in terms of asymptotic mass (*m*_*A*_) with the parameter *a* related to fundamental cell properties. The energy needed to create a cell, scales to the ¾ power as an individual grows; *a* is therefore equivalent to energy allocation rate, which is expected to vary with resource supply (see Table 1 for a list of MTE parameters). We recognise that variation does exist, for example the rate at which the exponent scales ^73^. Studies suggest it may be higher in some fish ^74^, therefore incorporating this may improve model accuracy, especially if such data were acquired for Atlantic salmon.

A classical sigmoidal shape, as seen in other individual growth models is derived by integrating equation (4), giving the post-embryonic growth model:

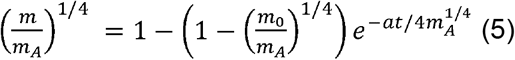

The temperature dependence of growth is also contained in parameter *a*, which expressed in terms of degrees Celsius, *T*_*c*_, is equal to 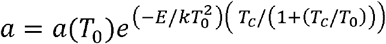 ^69^. Within *a, T*_0_ is a reference temperature equal to 273 K, as an arbitrary temperature at which water freezes and biological reactions cease ^69^. This can be rewritten as *a* = *a* (*T*_0_) *φ* (*T*_*c*_), where 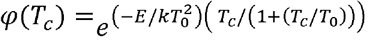. Substituted into equation (5) this provides a general expression of growth, relating development time to body mass and temperature:

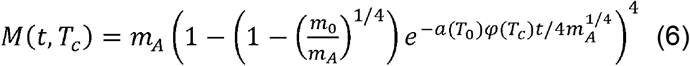

Here, the mass achieved starting from *m*_0_ where time *t* (in days), varies under different temperatures. A plot of equation (6) is exemplified at a range of temperatures across the natural logarithm of body mass in Fig. 1b.

To apply equation (6) to the growth kernel within the IPM, it can be rearranged to get the inverse function, with the time it takes for a fish to develop from day 0 as a function of mass:

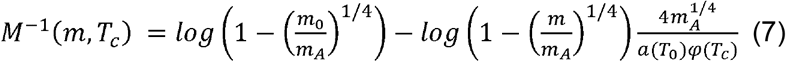

Equations (6) and (7) can then be used to calculate the mass reached after time, *t*, under temperature in degrees Celsius, *T*_*c*_, using

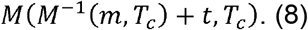

Here, *t* is equal to one day, as the smallest time scale to observe Atlantic salmon population dynamics. The variance in body mass at the next time step is assumed to follow a normal distribution, where the standard deviation is equal to the width of the cells in the IPM, *h*, calculated from dividing the length domain *m*_0_ to *m*_*A*_ into 300 small discrete bins. This results in the smallest variance there can be (as variance is unknown). The probability density function describing growth is therefore:

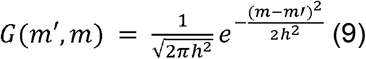

Note that the ontogenetic growth model applies to optimal (non-limiting) resource conditions, and accordingly, has largely been tested under idealised laboratory conditions. We therefore explore the effect of temperature on a population at a range of different values for the temperature-corrected normalisation constant of the growth equation, *a (T*_0_), to emulate varying resources. This is supported by *a* being later modified to incorporate resource supply ^75,76^. However, as we do not explicitly redefine *a (T*_0_) in terms of resource supply per se, for simplicity, we refer to it as “resource level” to refer to both the actual rate of resource availability as well as constraints imposed by the habitat on the ability of individuals to access those resources.

We derive a value for *a (T*_0_) from Atlantic salmon laboratory growth data, over a range of temperatures under non-limiting resources, collected from the literature (see Supplementary Methods). Here, rearranging equation (5), the growth model was fit to this data using

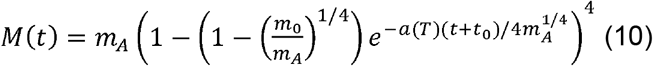

where *t*_0_ is an added constant to account for growth rates from a range of starting experimental masses. From this, the range of values for *a (T*) are plotted against 1/*kT* − 1/*kT*_0_, with the resulting intercept the metabolically optimal and temperature-independent value (see Table 1). We then test this initial value derived from laboratory data (termed metabolically optimal), along with a 25% and 50% reduction in this value to investigate how varying resource levels affect *r*_max_.

### Survival

The survival function *S*(*m*) is the survival probability from time *t* to *t* + 1 of an individual of mass *m*, and is also calculable from MTE. Individual-level metabolic processes are inextricably linked to population-level processes such as *r*_max_, as metabolism sets the demand for environmental resources and the resource allocation to vital rates^71^. Mortality within a population is therefore expected to have the same dependence on body mass and temperature ^22,28,29,77^. The relationship to temperature can be fitted with an inverse Sharpe-Schoolfield model ^22,78,79^. Despite this being a more prominent relationship in smaller size classes^80^, we apply this relationship to the whole size range of individuals with the assumption that smaller sizes are important for predicting population-level responses^81^, and the effect of a flattening unimodal relationship as individuals get larger would be negligible. Intrinsic mortality with an inverse Sharpe-Schoolfield temperature response for all individuals in a population is therefore be calculated by

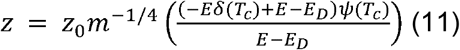

where 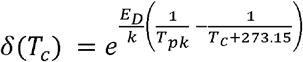 and 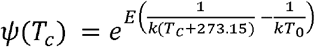 . Here, *z* is the instantaneous mortality rate and *z*_0_ is the mass- and temperature-independent normalisation constant. This equation is parameterised with data from brown trout (*Salmo trutta*) daily mortality data (see Supplementary Methods). Here, we derive estimates for *T*_*pk*_, the peak performance temperature (or in this case the inverse), *E*_*D*_, the deactivation energy and the normalisation constant, *x*_0_ (where *x*_0_ = *z*_0_ *m*^−1/4^). To correct for mass (*m* = 0.05; ^82^) we can derive an initial estimate for *z*_0_ from *x*_0_:

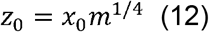

This however, results in a value for leading to a mortality rate that is higher than expected when compared to the mortality normalisation estimate from Savage et al. (2004)^71^, which utilises data across adult fish species and an Arrhenius temperature response. Despite Atlantic salmon and brown trout being more closely related than other salmonids ^83^, we recognise that there may be discrepancies in mortality rates here due to the embryonic life stage having a different physiology and therefore temperature tolerance. As a result, we shift this estimate of to align with MTE giving an estimate for natural mortality applicable across life stages (see Supplementary Methods). We compare our results to a higher mortality scenario through increasing the normalisation constant by 25%, as reduced survival may occur in the field due to factors such as increased predation or detrimental anthropological effects. However, we find that this has a lesser effect on *r*_max_ compared to varying resource level, or (see Supplementary Figure 1).

To incorporate mortality within the IPM, the probability of surviving in a day can be calculated from, equation (11), by taking 1 from the cumulative distribution function of an exponential distribution. Therefore, for any given mass the probability of surviving in a day is given by

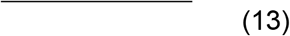

A plot of this equation at a range of temperatures (in degrees Celsius) across a range of body masses is exemplified in Fig. 1c.

### Reproduction

A measure of fecundity within salmonids is the clutch size; the number of eggs produced during spawning ^84^, and this is expected to increase with female body size ^85^. Within the fecundity kernel, the function considers the number of eggs produced by fecund life stages (specified by an adult maturation mass threshold,) to emulate a spawning event:

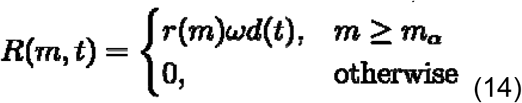

where is the number of eggs produced for mass and accounts for the sex ratio within populations, equal here to 1/2, halving the number of eggs produced to model the dynamics of females and assuming an equal sex ratio within populations ^86^. Finally, is the day of reproduction, equal to 1 when there is spawning day within the MIPM, otherwise this is equal to 0. To calculate the number of eggs produced across body mass (), we collated literature data on the number of eggs produced per female body size from a range of adult ages ^87–89^. As some body size measurements were reported as lengths, a body weight to length regression (from ^89^) was used to standardise all adult lengths into weights. The resulting allometric model for number of eggs produced is equal to

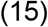

where is a constant calculated from the intercept and is the scaling exponent calculated from the slope of the data. A plot of this relationship is shown in Fig. 1d.

The size distribution of eggs produced per mature individual is expected to follow a normal distribution with a mean of 0.125g and standard deviation of 0.2 ^90–92^:

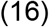

where *N* represents a normal distribution. The function *D*(*m*′, *m*) is then multiplied to the number of eggs produced (*R(m, t*)), giving the *F*(*m*′, *m,t*) kernel.

Temperature effects on Atlantic salmon fecundity were not incorporated here as there is not an evident correlation between temperature and the number of eggs produced in salmonids ^93^. Temperature is expected to most likely modify the spawning migrations and movements of salmonids rather than affect their physiology of the number of eggs laid ^94^, as well as mostly affect the incubation time before hatching ^95,96^. Temperature-dependent incubation time is however implicitly included within the model, through having a mass range within the egg stage, which experiences growth similarly to the other life stages within the MIPM. Thereby, under warmer temperatures the egg stage ‘hatches’ (or reaches the appropriate mass) quicker according to growth equation (6).

Within the MIPM, the *F*(*m*′, *m,t*) function is applied as a single daily event, as spawning is an instantaneous process in female Atlantic salmon ^97,98^. This takes place at the same time of year each year within the MIPM, as Atlantic salmon spawning events within regions are found to take place typically at the same time annually ^99^. However, the spawning day (where the reproduction function is applied within the kernel) is an adjustable parameter in the model, as this spawning date may vary across the Atlantic salmon’s range ^100^. The nature of the model then allows egg hatching and progression to the next stage to be dependent on temperature, which is shown to be more variable than spawning date among salmonids and is influenced by temperature ^100^.

### Calculating population fitness

To initialise the system, an abundance vector of 1000 eggs were added with a burn in time of 40 years. This allows the modelled populations to stabilise; a feature of structured population models ^86^. Subsequent analyses then only used data collected after the burn in period. For mean temperature scenarios, a range of fixed temperatures from 0°C to 30°C were modelled for 30 years, with 365 days in a year, at the contrasting parameter values for *a (T*_0_). Fluctuating seasonal temperature scenarios were modelled for the same time frame using a sine function, with a decreasing amplitude as mean annual temperature increases. This is due to seasonality varying geographically and with a predictable relationship to temperature, in that regions with a higher mean annual temperature also have a less variable temperature regime across months ^108–110^. We therefore derived a negative linear relationship between amplitude and mean annual temperature from time series temperature data from four known Atlantic salmon rivers to parameterise a sine function, reflecting real-world gradients (see Supplementary Methods).

The application of a projection matrix within IPMs enables population growth rates (the dominant eigenvalue) to be easily calculated (after discretisation of the mass variable), to investigate population fitness ^86^. We therefore calculated the maximal rate of population growth (*r*_max_, where *r*_max_ = log(*λ*)). Negative *r*_max_ values indicate population decline and positive ones, growth, therefore *r*_max_ = 0 results in a population remaining stable, but in practice *r*_max_ >0 would lead to stable populations after some density dependent factors take effect that are not currently captured in the model. We then compared *r*_max_ to the natural thermal environment that extant Atlantic salmon are found in (using^18,111–116^), giving the empirical realised thermal niche.

Specifically, to characterise this, we extensively searched the literature for mean annual river temperatures from northern populations in Norway and Canada to southern populations in Spain. This gives a mean annual freshwater temperature range of 3°C to 15°C. We explored the effect that mean global warming scenarios will have on the empirical realised thermal niche, assuming populations do not adapt or shift their distribution as a response. We assumed that the extant range of Atlantic salmon would therefore increase in temperature under an IPCC scenario of 1.5°C (SSP1-1.9^55^) and 3.6°C (SSP3-7.0^55^).

### Signatures of resource level and temperature on the size distribution

The mean size distribution from August in the time series was calculated across all mass divisions and the median mass of this size distribution determined. This was calculated from the mean abundance for every mass division during August and the subsequent proportions compared. The median mass from the size distribution was measured in August as (i) typically population survey/monitoring data is collected during the summer months ^11^, and (ii) it allows enough time to have passed from egg laying in the winter months to observe a cohort’s progression on the size distribution (a cohort is defined as all members of a population produced over a single reproductive event^117^). We excluded the median mass from plotting where *r*_max_ is negative, as the mean size distribution is not meaningful as a population goes towards extinction.

## Data and code availability

All data and code for reproducing the study’s analyses can be found at https://github.com/oliviamorris/MIPM

## Acknowledgements

We thank Six Rivers Iceland for funding this work set out to explore declines in Icelandic populations of Atlantic salmon, as a PhD project to OM. We would also like to acknowledge helpful discussions and interactions with Peter Williams, Tom Clegg, Paul Huxley and scientists at the Marine and Freshwater Research Institute, Iceland. Finally, AGF would like to acknowledge financial support from the Government of Cantabria through the Fénix Programme.

## Notes

### Competing Interest Statement

The authors have declared no competing interest.

