## Supplementary Figures for "Climatic limits to Atlantic salmon population fitness at continental scales"

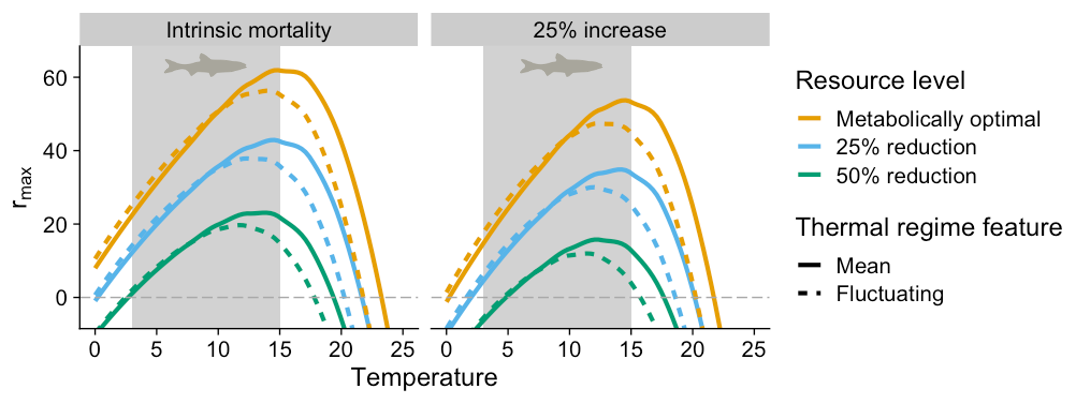

**Supplementary Figure 1.** **The effect of differing thermal regimes and resource level on population fitness, *r*_max_ with reduced survival.** The effect of increasing mean and seasonal fluctuating temperatures (°C) on the maximal rate of population growth, *r*_max_. A 25% decrease in survival for Atlantic salmon has less of an effect than varying resource levels.

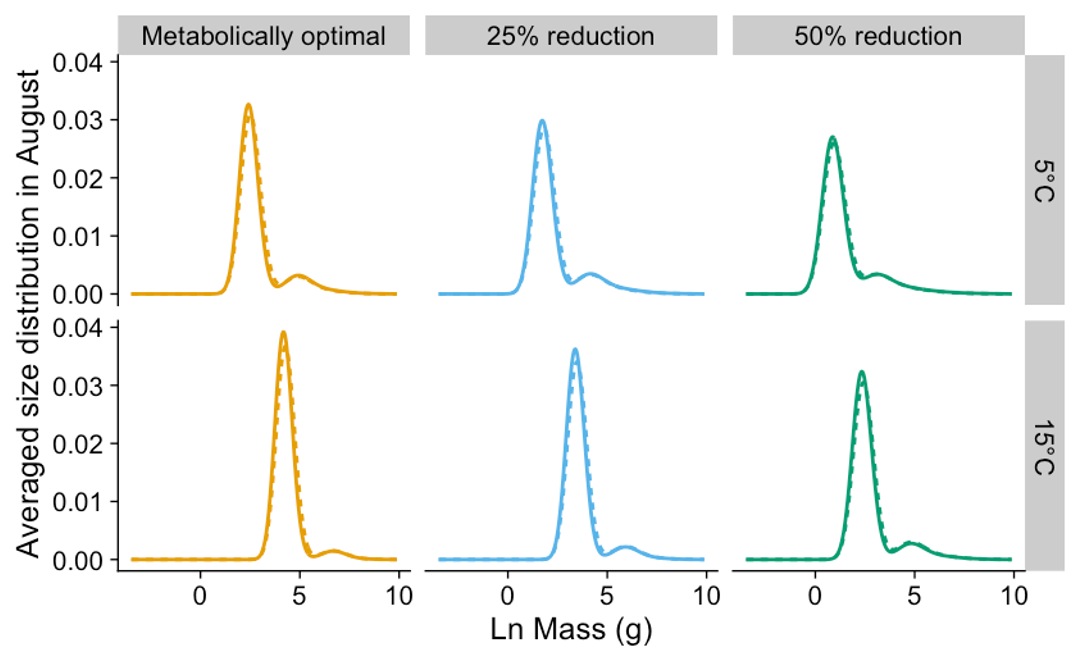

**Supplementary Figure 2.** **The size distribution averaged during August across resource level.** The effect of a mean of 5 and 15°C and seasonal fluctuating temperatures with a mean of 5 and 15°C on the averaged size distribution during August for Atlantic salmon from the MIPM for mean (solid line) and fluctuating (dotted line) thermal regimes. Note that this is the size distribution during August as this is typically when empirical sampling takes place (see Methods).
