## Supplementary Methods for "Climatic limits to Atlantic salmon population fitness at continental scales"

### Growth

We derive a value for $a(T_{0})$ from Atlantic salmon laboratory growth data, over a range of temperatures under non-limiting resources, collected from the literature ^1–3^ (Supplementary Methods Fig. 1a). Equation 10 (Methods) was fit to this data and from this, the range of values for $a$ are plotted against $1/kT - 1/KT_{0}$, with the resulting intercept the metabolically optimal and temperature-independent value $a(T_{0})$ (Supplementary Methods Fig. 1b).

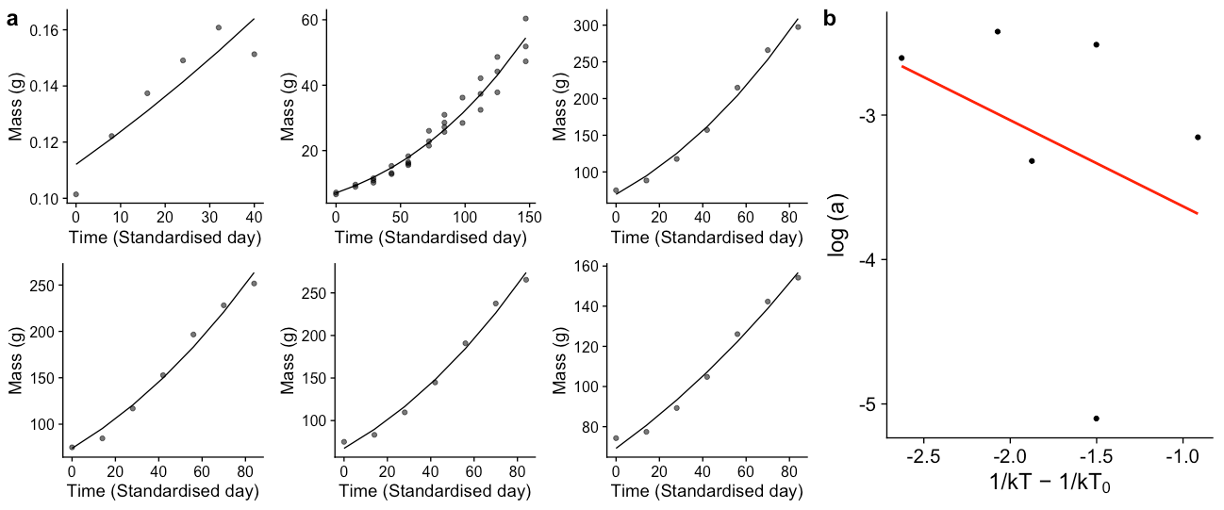

**Supplementary Methods Figure 1. Metabolically optimal parameter derivations for individual growth rates.** An initial parameter estimate for $a(T_{0})$ was derived from individual experimental growth rate data under non-limiting resources (a). Fitting the growth model (Methods equation 10) estimates for $a$ were derived. These were then plotted against the inverse temperature from which the studies were collected to get an initial value for temperature-independent energy allocation ($a(T_{0})$) from the intercept of a linear model (in red) fitted (b).

### Survival

The normalisation constant for survival ($z_{0}$) results in a value for mass- and temperature-independent $z_{0}$being too high compared to Savage et al. (2004)^4^. We therefore compare the unimodal relationship of survival to the Arrhenius relationship, and shift mortality to line up with this (Supplementary Methods Fig. 2).

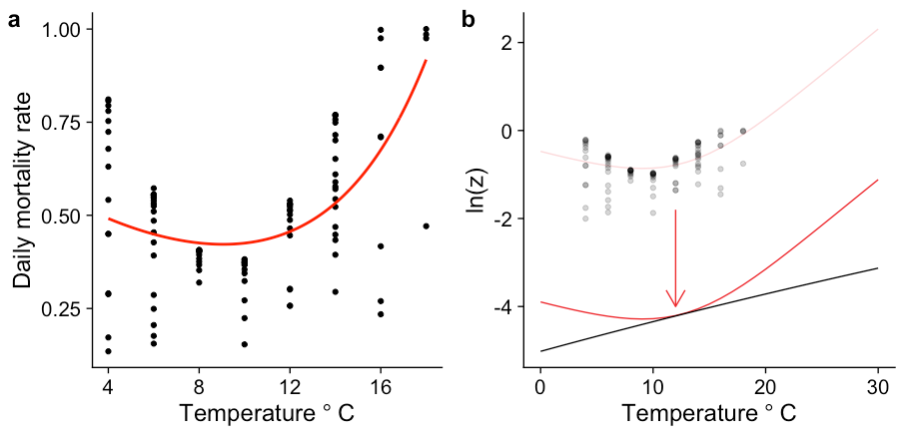

**Supplementary Methods Figure 2. Survival model parameter derivations.** Initial parameter estimates were calculated from fitting the mortality equation 11 (Methods) to experimental laboratory data from brown trout (*Salmo trutta*) (a). The resulting model (in red) led to estimates for the peak performance temperature ($T_{pk}$), the deactivation energy ($E_{D}$) as well as an estimate for $x_{0}$ which when mass-corrected (from the mean mass of the measured individuals) arrives at an initial parameter estimate for $z_{0}$ that is higher than expected. We adjusted the normalisation constant (b) to align with the Arrhenius mortality relationship parameterised with adult data across fish species from Savage et al. (2004)^4^ (black line).

### Fluctuating seasonal temperature derivation

Seasonal variation varies predictably with mean annual temperatures. We therefore estimate this from four known Atlantic salmon rivers across species range. We derived a relationship between amplitude and mean annual temperature from time series temperature data from four known Atlantic salmon rivers to parameterise a sine function, reflecting real-world gradients. Where there was missing data in the time series due to sampling error, this was imputed with overall daily mean values. Then, using non-linear least squares regression, the amplitude was estimated across each location. These were plotted against mean annual temperature for each location, giving a negative linear relationship ($y=-0.02x+5.43$), relating increasing mean temperatures to amplitude for northern hemisphere Atlantic salmon rivers (Supplementary Methods Fig. 3).

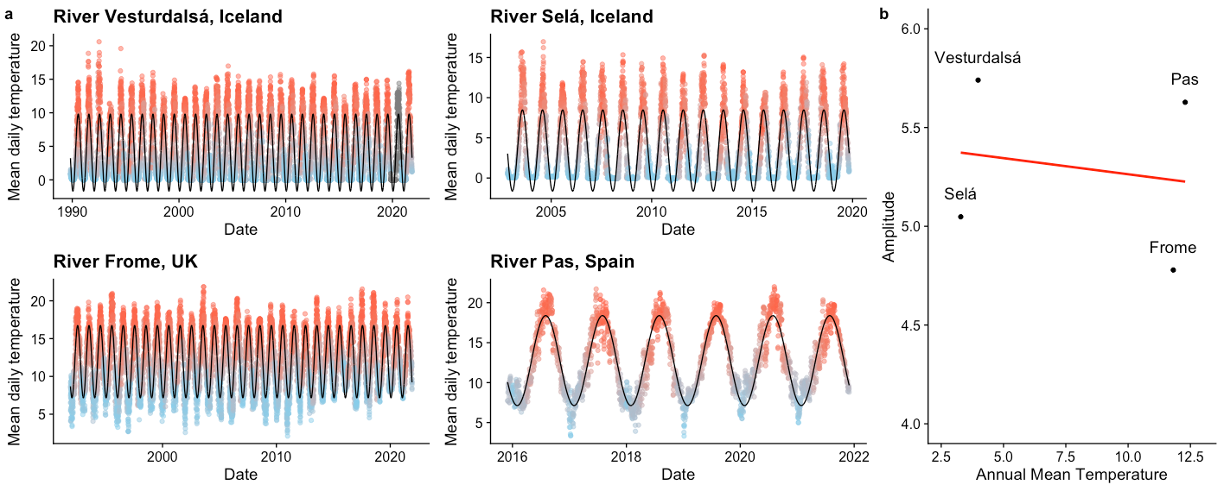

**Supplementary Methods Figure 3. Seasonal temperature fluctuations calculated from amplitudes within Atlantic salmon river temperature data.** Within-river mean daily temperatures from known Atlantic salmon rivers spanning the Atlantic salmon's distribution (a), including the Vesturdalsá, Iceland, the Selá, Iceland, the Frome, UK and Pas, Spain. Imputed daily temperatures calculated from the time series mean are shown as grey dots, with each location then fitted with a sine function to calculate the amplitude (black line). The resulting amplitude values are fitted against the annual mean temperature for each location (b) to get a linear relationship (red line) of which to calculate how amplitude may change across the range of climatic regimes Atlantic salmon span.
